# Cellular Orientation Is Guided by Strain Gradients

**DOI:** 10.1101/095976

**Authors:** Sophie Chagnon-Lessard, Hubert Jean-Ruel, Michel Godin, Andrew E. Pelling

**Affiliations:** Department of Physics, Center for Interdisciplinary Nanophysics, 598 King Edward, University of Ottawa, Ottawa, ON, K1N 6N5, Canada., (6965); Department of Electronics, 1125 Colonel By Drive, Carleton University, Ottawa, ON, K1S 5B6; Department of Mechanical Engineering, Site Building, 800 King Edward Avenue, University of Ottawa, Ottawa, ON, K1N 6N5, Canada; Ottawa-Carleton Institute for Biomedical Engineering, Ottawa, Ontario, K1N 6N5, Canada; Department of Biology, Gendron Hall, 30 Marie Curie, University of Ottawa, Ottawa, ON, K1N 6N5, Canada; Institute for Science, Society and Policy, Desmarais Building, 55 Laurier Ave. East, University of Ottawa, Ottawa, ON, K1N 6N5, Canada

## Abstract

The strain-induced reorientation response of cyclically stretched cells has been well characterized in uniform strain fields. In the present study, we comprehensively analyse the behaviour of human fibroblasts subjected to a highly non-uniform strain field within a polymethylsiloxane microdevice. Our results indicate that the strain gradient amplitude and direction regulate cell reorientation through a coordinated gradient avoidance response. We provide critical evidence that strain gradient is a key physical cue that can guide cell organization. Specifically, our work suggests that cells are able to pinpoint the location under the cell of multiple physical cues and integrate this information (strain and strain gradient amplitudes and directions), resulting in a coordinated response. To gain insight into the underlying mechanosensing processes, we studied focal adhesion reorganization and the effect of modulating myosin-II contractility. The extracted focal adhesion orientation distributions are similar to those obtained for the cell bodies, and their density is increased by the presence of stretching forces. Moreover, it was found that the myosin-II activity promoter calyculin-A has little effect on the cellular response, while the inhibitor blebbistatin suppresses cell and focal adhesion alignment and reduces focal adhesion density. These results confirm that similar internal structures involved in sensing and responding to strain direction and amplitude are also key players in strain gradient mechanosensing and avoidance.

## Insight, innovation, integration

In the body, mechanical stress arising due to movement exposes cells to complex and anisotropic strains and strain gradients. Employing an innovative microfabricated device, we have uncovered how strain gradients can act as an important biological signal. Our device enables the systematic investigation of strain and strain gradient directions in a single membrane. Decoupling these two pieces of mechanical information provides critical new evidence that cells are able to spatially integrate and respond to both physical cues. Moreover, cells specifically respond to these two simultaneous physical cues by exhibiting a clear strain gradient avoidance behaviour. This work reveals new insights into how strain gradients play a key role in guiding the long-range organization in populations of living cells.

## Introduction

A rapidly growing body of evidence has established that mechanical forces and force gradients can act as key drivers of biological processes at the molecular, cellular, and organismal scales^1–6^. The web of interactions between the cell and its microenvironment involves a complex interplay between mechanical forces and biochemical signals. These interactions are made possible by the cell’s ability to sense external mechanical cues, transduce them into intracellular biochemical signals, and generate short and long term responses that orchestrate crucial cellular functions. Such mechanisms have been reported to be involved in the regulation of a variety of cellular functions such as cell migration, division, apoptosis, differentiation, and gene expression^7^.

Another relevant example, which has become a case study for mechanosensitivity and response, is cell reorientation under cyclic stretching. Such stretching is experienced by cells in vivo during many processes including the pumping of the heart, the contraction of the muscles, and the expansion of the lungs. Through tissue matrix deformations and cytoskeleton-adhesion interfaces, many cell types are thus cyclically stretched, including human fibroblast^8^. It has been shown in numerous studies that adherent cells subject to cyclic uniaxial strain via a deformable extracellular matrix reorient themselves approximately perpendicularly to the strain direction to a degree that depends on the strain amplitude^9–19^. This was observed for frequencies above ∼0.1 Hz, although the characteristic time of the response was found to be frequency dependent^10^. Over the last few decades, different models and refinements have been proposed to describe cellular reorientation and explain the underlying mechanisms behind this phenomenon^11,12,20–24^. It is also well established that the cytoskeletal and focal adhesion (FA) dynamics are central to this process, as they form the contractile and mechanosensing machinery. Specifically, the disruption of contractile activity can lead to the inhibition of cellular alignment^25–27^. Nevertheless, a complete understanding of the mechanism by which applied physical forces influence the FA structures and the cytoskeletal remodelling, individually and as concerted dynamics, has yet to be reached^28–31^.

Simple strain fields can be approximated as being uniform, i.e. with a constant strain amplitude as well as a non-varying strain direction. In contrast, in complex non-uniform strain fields, the strain amplitude and potentially the strain direction vary spatially. This gives rise to a strain gradient field, also described by a specific gradient amplitude and gradient direction at each spatial location. In the majority of cases, advancements in the field have been achieved by using simplified substrate strain fields^10–12,14–17,19^. More recently, various other studies have employed ingeniously designed macro-devices with stretchable membranes that generate non-uniform strain amplitudes, both in the context of static and cycling stretching^9,32–40^. In a few cases^9,34,38^, this approach allowed for the successful analysis of cell reorientation (among other responses) as a function of the strain amplitude over a single non-uniformly stretched membrane. In most cases, the strain field is examined with reference to the average principal strain direction, taken as the x- or y-axis, therefore ignoring local variations in strain direction. Moreover, the strain gradients that arise in non-uniform and complex strain fields are rarely considered. Importantly, in all of these studies, strain gradient dependence was not reported in the context of cell reorientation.

While simple systems generating uniaxial stretching have the benefit of isolating specific mechanical cues, they do not reproduce the complexity and the non-uniformity of the strain fields occurring in vivo^33,41,42^. Moreover, the strain gradients in the body span multiple orders of magnitude (∼10°-10^2^ % mm^-1^)^36^ and are oriented along multiple directions with respect to the principal strain. Importantly, it is known that cells are able to sense chemical gradients^43^ as well as substrate stiffness gradients^44^, which were both demonstrated in the context of preferential cell migration. Therefore, we hypothesize that the strain gradient plays a key role in the phenomenon of cellular reorientation in non-uniform strain fields. Such a finding would imply that cells are able to sense multiple components of a complex strain field (amplitude and direction of the principle strain and strain gradient) and integrate these diverse mechanical cues in their response.

Here, we present a polydimethylsiloxane (PDMS) stretching microdevice (an earlier version of which was reported previously ^13^) that allows for the generation of a highly non-uniform strain field across the membrane surface, non-uniform principal strain directions, and non-uniform gradients. The necessity for a microdevice arises from the need for generating significant strain gradient amplitudes on the cellular scale while maintaining a maximum strain amplitude of approximately 10%. This falls in the amplitude range known to induce cell alignment following cyclic stretching^11,12,20^. Therefore, the design of our microdevice allows us to mimic complex in vivo physical forces that can play a critical role in cell biology.

Comprehensive analysis of human foreskin fibroblast (HFF) cell reorientation under cyclic stretching in the device employed in this study indicates that the final orientation of the cells is determined by both the strain- and the strain-gradient fields experienced by the cells individually. We found a strong correlation between the strain amplitude and the degree of cell normal alignment with respect to the principal strain direction (i.e. the degree to which the elongation axis of the cells aligns perpendicularly to the principal strain direction). More importantly, we were able to clearly demonstrate a similar correlation between the strain gradient amplitude and the degree of cell normal alignment with respect to the maximum principal strain gradient direction. To further investigate this cellular behaviour under cyclic non-uniform anisotropic strain, we examined the influence of the myosin-II activity on the cell reorientation as well as on the FA density and orientation. Under our experimental conditions, we discovered that the effect of the stretching force alone leads to a slight increase in FA density, while their orientation follows the direction of cell elongation. We also show that myosin-II contractility is a major player of the phenomenon of cell sensitivity to strain gradients arising in complex cyclic strain fields.

## Results

### Description of the microdevice’s non-uniform anisotropic strain

In order to produce biologically relevant strain gradients with directions that are decoupled from those of the strains, we developed a PDMS stretching microdevice. A schematic of this device is presented in Fig. S1 (ESI†), and details of its fabrication, working principle, and thorough characterization are provided in the Methods and Materials section and in the ESI†. At any position on the membrane, the direction and the amplitude of the maximum principal strain **ε_1_**(x,y) (maximum stretch) were extracted. The non-uniformity of the maximum principal strain gives rise to a gradient, ∇ε_1_(x,y), which can also be described by an amplitude and a direction. This amplitude represents the local rate of change of the maximum principal strain amplitude, and its direction points toward that of greatest change. Fig. 1 provides a visual guide to the key parameters involved in the cell orientation analysis. The underlying color map of Fig. 1A is that of the maximum principal strain amplitude profile ε_1_(x,y). In Fig. 1B, the lengths and directions of the blue lines represent the maximum principal strain vector **ε_1_**(x,y), amplitudes and directions respectively (Fig. 1B). ε_1_ is maximal near the top and bottom vacuum pumps (not shown), and is relatively constant in the central region. The red lines represent the amplitudes and directions of the gradient ∇ε_1_(x,y). The gradient directions are orthogonal to the lines of strain equi-amplitude (color lines in the amplitude map). It should be noted that the effect of the gradient on the cellular reorientation is most likely to be observed where the gradient amplitude is large while the strain amplitude is low, such as near the corners.

**Fig. 1.**
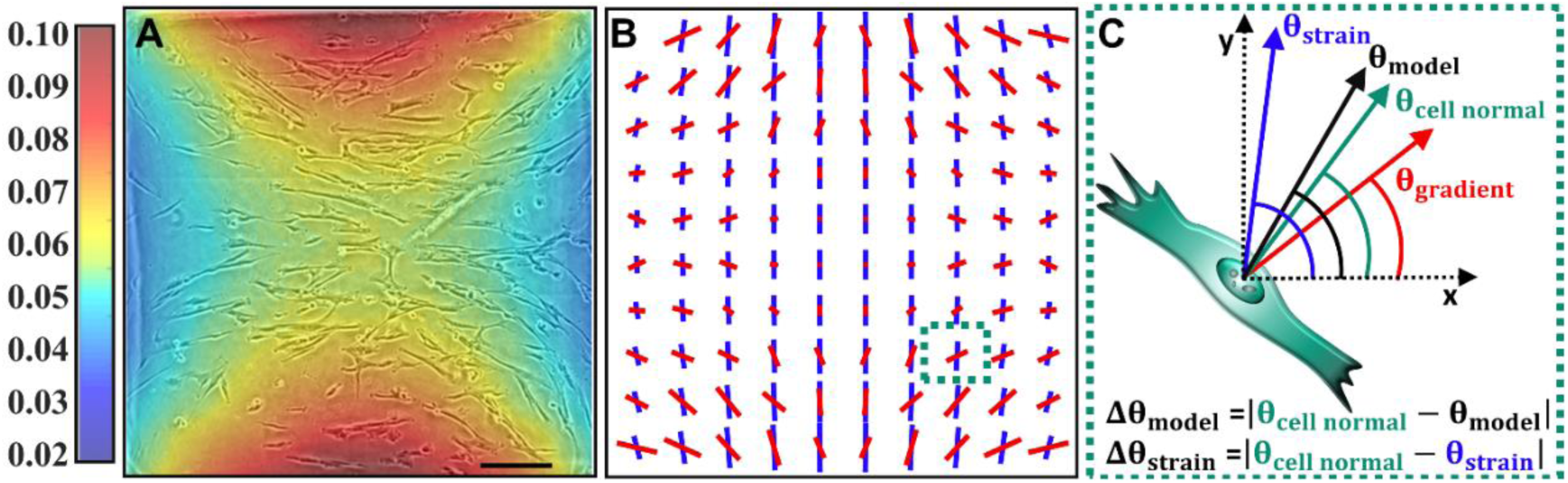
Description of the non-uniform anisotropic strain field and of the angle definitions. (*A*) Color map of the experimentally determined maximum principal strain amplitude ε _1_(x,y) across the 1.6 mm wide square membrane. A phase contrast image of adhered HFF cells on the microdevice membrane is overlaid on the strain map. The cells have been subjected to 11 hours of vertical cyclic stretching at 1 Hz. They do not simply reorient away from the strain direction (mainly vertical as seen in B) but they rather largely follow lines of equal strain amplitude. Note that the presented area is cropped slightly to exclude the potential edge effects (8% wide on each side). The scale bar is 350 μm. (*B*) A representation of the amplitude and direction of the maximum principal strain (blue lines) and its gradient (red lines) is displayed. (*C*) Scheme of a fictional cell located in the bottom right corner of the device (dotted square shown in B) presenting the definitions of the angles used in the analysis. The angles Δθ_model_ and Δθ_strain_ are used to construct the reorientation histograms. Note that the relative weight of θ_gradient_ and θ_strain_ to obtain θ_model_ depends on their relative amplitude, as explained in the main text.

The complex strain field generated by this device enables the differentiation between the cellular response induced by the strain field and that induced by the strain gradient field. Many previous studies have employed a largely uniform and anisotropic strain, which allowed them to report the data on a fixed axis^11,12,14–17,19^. In our case, the non-uniformity of the field requires a different analysis method. Since the maximum principal strain direction varies, the cell orientation angle with respect to a single arbitrary axis are inappropriate if cells from different locations on the membrane are to be included in the same analysis. This is especially important when the gradient direction is considered in the description of cell orientation. In the sections that follow, the orientation of the cell normal is computed against either the local maximum strain orientation or against an empirical model that we have developed that considers both the local strain and its gradient. The definitions of the different angles used in the analysis are summarized in Fig. 1C.

### Cell reorientation depends on both the strain and the strain gradient fields

Multiple experiments were performed to investigate the reorientation behaviour of HFF cells. Fig. 1A shows a representative image of adhered cells on the membrane embedded in the microdevice after 11 hours of cyclic stretching with the non-uniform strain field. The cells were initially randomly oriented (not shown), and after cyclic stretching they were approximately perpendicular to the maximum strain direction in the region of constant strain (central region). Interestingly, outside of the central region, the cell orientations approximately followed the underlying color map, which represents the principal strain amplitude profile. More specifically, they appeared to align preferentially along lines of equal strain amplitude. In other words, they seemed aligned perpendicularly to the strain gradient direction, especially in the region of strong gradient amplitude (rapid change of color).

Fig. 2A-C (excluding the inset of Fig. 2A) present the histograms of the angle differences between the cell normal directions and the principal strain directions (Δθ_strain_) considering three different regions of the membrane. The full membrane surface is examined in 2A while only the regions of high strain amplitude (> 7%) are analyzed in 2B, which correspond to the vacuum pump neighbouring regions (orange and red regions in Fig. 1A). The histogram of Fig. 2A confirms that the cells largely reoriented perpendicularly to the strain direction and the larger degree of alignment observed in 2B shows that this reorientation is sensitive to the strain amplitude. In contrast, Fig. 2C considers regions of simultaneously high gradient amplitude (> 7% mm^-1^) and low strain amplitude (< 5%) (green and turquoise regions near the corners of Fig. 1A). Under these conditions, a decrease in cell normal alignment with the strain direction is observed, suggesting that this description is incomplete. In particular, in these regions, the cell normal directions seem to depart from the principal strain directions and align partly with the gradient directions. It should be noted that in our device, the principal strain and gradient directions are largely decoupled, allowing this observation.

**Fig. 2.**
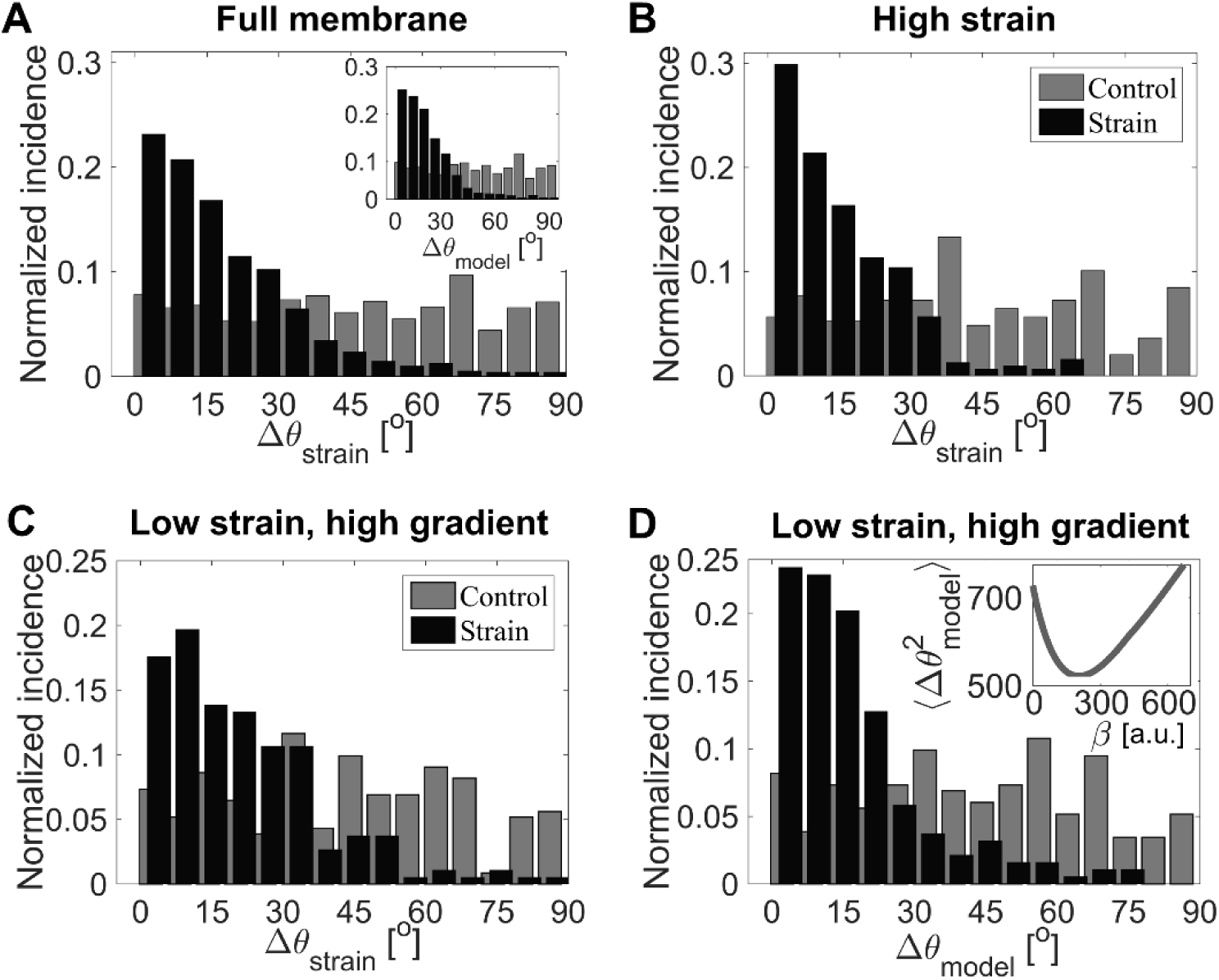
Reorientation analysis of the HFF cells after 11 hours of cyclic stretching. Normalized incidence histograms of Δθ_strain_ (*A*-*C*) and Δθ_model_ (*D*, inset of *A*) for both unstretched controls and stretched experiments. All the cells are included in (*A*) (main figure and inset), only the cells located in the regions of high strain amplitude (> 7%) are included in (*B*), and only cells located in the regions of simultaneously high gradient amplitude (> 7% mm-1) and low strain amplitude (<5%) are included in (*C*, *D*). It should be emphasized that for all the histograms, the cell normal angles are not reported with respect to the fixed x-axis, but rather with respect to the underlying maximum principal strain directions or to the empirical model directions, as specified. The inset of (*D*) shows the optimization curve of the β value via mean square minimization of the Δθ_model_ values. Each histogram comprises the combination of 6 unstretched control experiments as well as the combination of 6 cyclically stretched experiments. The number of cells analysed in each histogram is n_strain_=1078 and n_control_=1244 for (A), n_strain_=132 and n_control_=200 for (B), n_strain_=181 and n_control_=187 for (C, D). Fig. S3 of the ESI† explicitly shows the different regions used to generate Fig. 2A-D, and table S1 summarizes the number of cells analysed in each case for both the control and stretched experiments.

In an attempt to describe the cell reorientation under such a complex strain field, a simple empirical model was used. Building on our results and recurrent observation that cells seem to align largely with equal strain amplitude lines, we incorporate the following ideas: i) cellular avoidance of stretching (strain) in regions of strong applied strain amplitude and ii) cellular avoidance of strain gradient in regions of strong applied gradient amplitude (identified in this paper). To take into account these effects, we consider the simplest empirical model possible, namely a linear combination of θ_strain_(x, y) and θ_gradient_(x, y) weighted by the strain amplitude ε_1_(x, y) and the strain gradient amplitudes |∇ε_1_(x, y) |, respectively. To investigate our data, the following equation is thus suggested to express the preferential orientation of the cell normals:

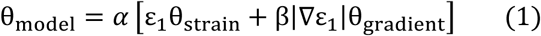

where *α* = (*β*|∇ε_1_| + ε_1_)^-1^ is the normalization factor including the weighting parameter β. Fig. 2D presents the histogram of the angle differences Δθ_model_ (differences between θ_Cell normal_(x, y) and θ_model_(x, y)), considering again the regions of high gradient and low strain amplitudes (same areas as in 2C). The parameter β was optimized here (and kept constant for all subsequent analysis) by minimizing the mean squares 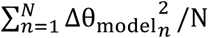 considering the data from six distinct stretching experiments. Importantly, the optimal β value remains similar (within ∼ 13%) whether we consider the full membrane or the regions of interest in 2D. Interestingly, while the model provides a slight improvement over the full membrane (Fig. 2A inset), it is not comparable to the excellent improvement obtained from Fig. 2C to Fig. 2D. The reason is that the gradient is relatively low over a large portion of the membrane, where the strain is thus dominant, resulting in a dilution of the gradient effect. In the regions where the gradient effect is most important (Fig. 2C, D), the experimental and “predicted” orientations are in good agreement only when the model is considered. It is interesting to note that for the regions considered in Fig. 2D, the average contributions of the strain and strain gradient (considering the optimal β value) are respectively 63% and 37%. This confirms the significance of the effect of the strain gradient on the cellular reorientation in our strain field. Finally, for Fig. 2A-D, the controls were randomly oriented.

### Cell alignment with the gradient direction depends on the gradient amplitude

We now further assess the dependence of the cell reorientation on the strain gradient in order to test its accuracy. First, Fig. 3A indicates that as the strain amplitude ε_1_(x, y) increases, the mean cell normal direction increasingly aligns with the principal strain direction (showed by a decreasing Δθ_strain_value). A similar relationship for the strain gradient is successfully reported in Fig. 3B. It indicates that as the strain gradient amplitude |∇ε_1_(x, y)| increases, the mean cell normal direction increasingly aligns with the gradient direction (showed by a decreasing Δθ_gradient_ value). To verify that the latter dependency cannot be attributed to a correlation between the strain and the strain gradient fields, two additional relationships were extracted. First, the mean Δθ_strain_ was plotted as a function of the gradient amplitude|∇ε_1_(x, y)| (Fig. 3C). Second, the average angle difference between the strain and the gradient directions, < θ_strain_ – θ_gradient_ >, was also plotted as a function of the gradient amplitude (Fig. 3C inset). In both cases, the unambiguous trend observed in Fig. 3B is not reproduced, indicating the genuine role of the gradient. In other words, the increased alignment of the cell normal with the strain gradient direction observed in Fig. 3B cannot be explained by the relationship between θ_strain_ and θ_gradient_. Importantly, for strain gradient amplitudes greater than ∼2% mm^-1^, the mean Δθ_gradient_ in Fig. 3B is systematically lower (“better”) than < θ_strain_ – θ_gradient_ > (Fig. 3C inset). To our knowledge, this is the first reported demonstration of cell alignment with the underlying strain gradient.

**Fig. 3.**
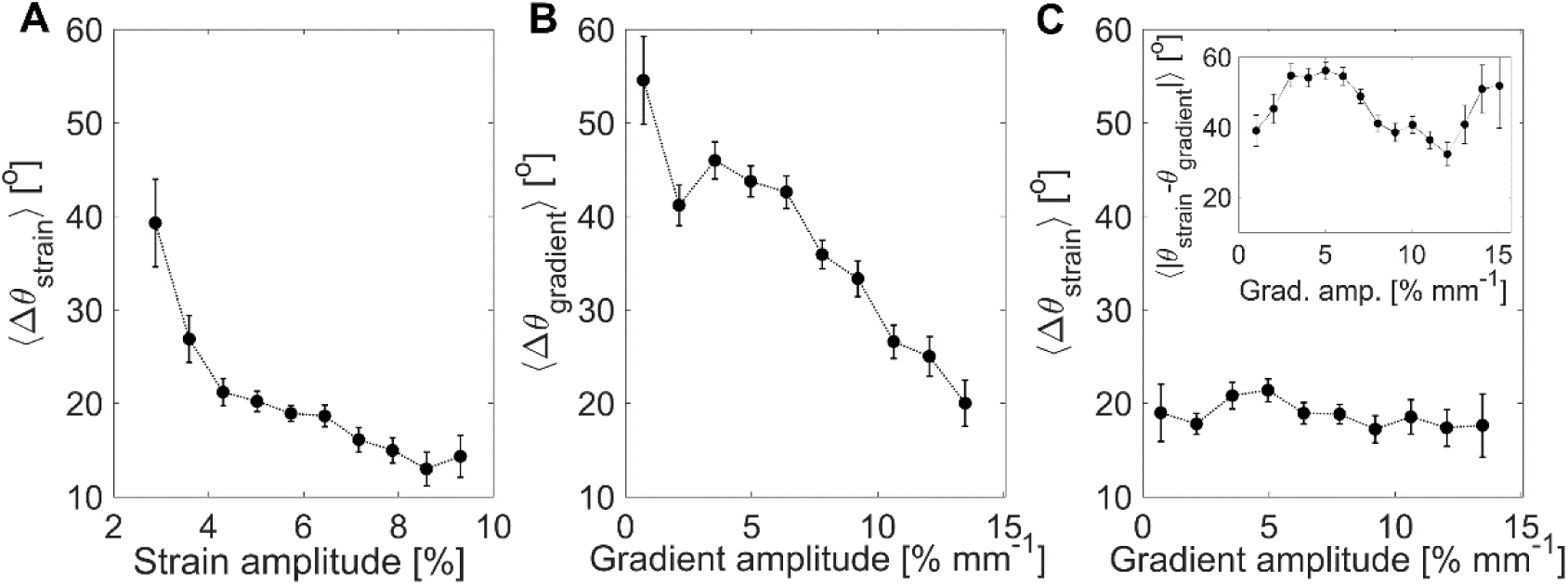
Dependence of the cell reorientation direction on the strain amplitude and strain gradient amplitude. The data were binned in equal amplitude range sizes and the error bars represent the standard error of the mean within each bin. (*A*) Mean angle difference between the experimental cell normal orientation and the principal strain direction as a function of the strain amplitude. (*B*) Mean angle difference between the experimental cell normal orientation and the strain gradient direction as a function of the strain gradient amplitude. (*C*) Mean angle difference between the experimental cell normal orientation and the principal strain direction as a function of the strain gradient amplitude. The inset shows the average angle difference between the strain and the gradient directions as a function of the gradient amplitude. The number of cells analysed is n=1078.

### Myosin-II activity inhibition supresses cell alignment while its promotion does not impact the final reorientation

To assess how myosin-II activity affects the orientation response of HFF cells in a non-uniformly strained microenvironment, experiments were performed in which calA or blebbistatin treatments were applied immediately before activating the substrate cyclic stretching ^27,45,46^. Fig. 4A shows that upon stimulation of myosin-II contractility through inhibition of myosin light chain phosphatase (calA), the cells reoriented in a similar fashion to their untreated counterparts (Fig. 2A, inset), with no significant difference. Conversely, the inhibition of myosin-II contractility largely prevented stretch-induced alignment under cyclic stretching, as shown by the near-random distribution of cell orientations displayed in Fig. 4B. It should be reiterated that the weighting parameter β employed in the orientation analysis of Fig. 4 was not re-optimized. However, it is very interesting to note that an optimization of the β value on the calA-treated cells of Fig. 4A would result in a β value that defers by only ∼ 8%.

**Fig. 4.**
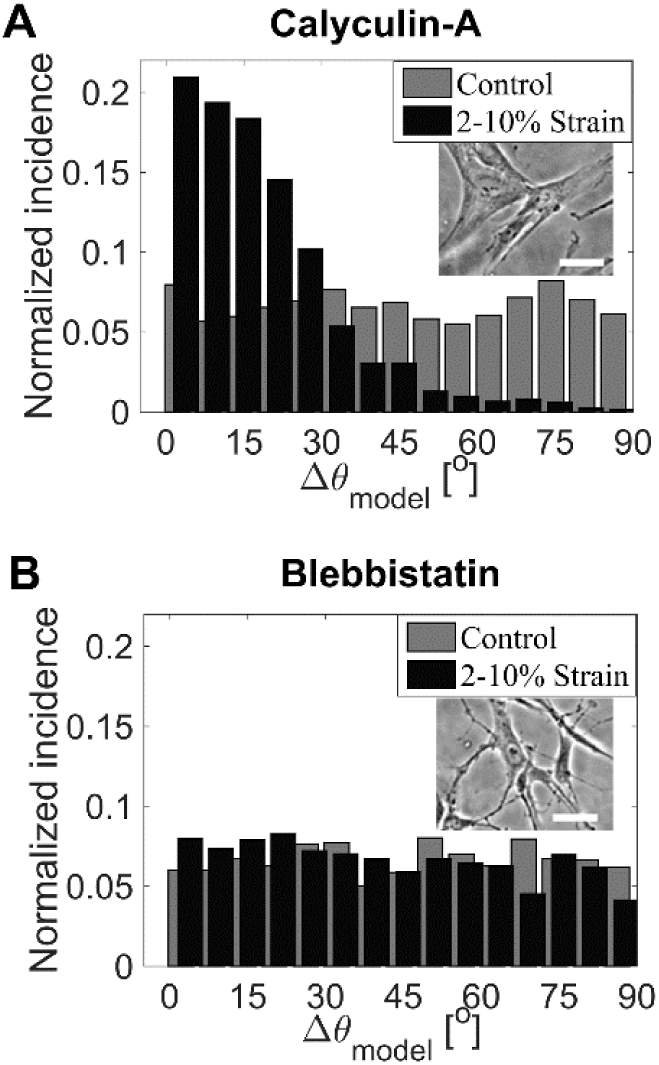
Effect of myosin-II activity on HFF reorientation after 11 hours of cyclic stretching. Normalized incidence histograms of Δθ_model_ for cells treated with (*A*) calA (2 nM) and (*B*) blebbistatin (10 μM). Each histogram comprises the combination of 6 unstretched control experiments as well as the combination of 6 cyclically stretched experiments. The number of cells analysed in each histogram is n_strain_=1231 and n_control_=1255 for (A), and n_strain_=1319 and n_control_=1091 for (B). The insets diplay phase constrast images of representative cells from the corresponding experiments. Scale bars are 100 μm.

### Focal adhesions reorient along the cell elongation direction under cyclic stretching and their reorganization is myosin-II dependent

To gain insight on the cellular reorientation mechanism, the behavior of the FA complexes was also assessed since they sense and transmit to the cell the physical signals from the extracellular matrix. Fluorescence images of a representative stretch-induced reoriented cell inside a microdevice are displayed in Fig. 5E-F. It can be seen that the actin filaments were largely aligned with the cell elongation direction, as expected ^17,20,30^. This qualitative observation applied to all untreated and calA-treated cells, but not to those treated with blebbistatin, in which case the alignment was less obvious. The FA protein vinculin, shown in Fig. 5F, also exhibited strong alignment with the cell direction. This is quantitatively assessed in Fig. 5A, in which the FA normals are seen to predominantly follow θ_model_ directions, in accordance with the cell elongation directions (Fig. 2A, inset). An equivalent FA orientation behaviour was observed for the cells with increased myosin-II contraction (calA, Fig. 5B), but little preferential alignment was found in the case of those with decreased contractility (blebbistatin, Fig. 5C). Interestingly, this is also reflected in the FA density (Fig. 5D). In comparison to the untreated cells, there was a significant loss of FAs in blebbistatin-treated cells, but not in those treated with calA. Finally, in all cases, the FA density was slightly strain-dependent; more specifically the applied stretching forces appeared to promote FA assembly.

**Fig. 5.**
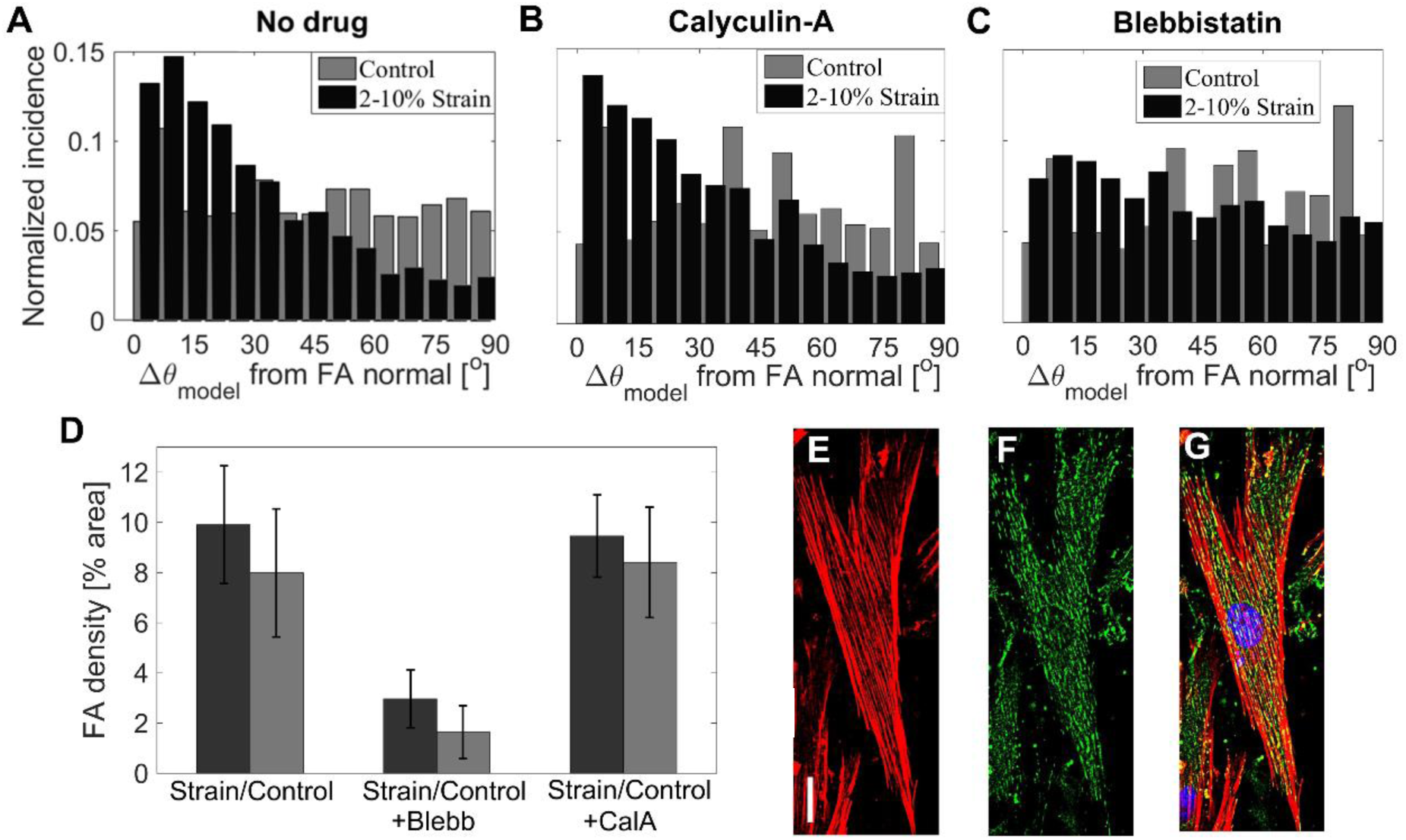
**FA reorganization after 11 hours of cyclic stretching.** Normalized incidence histograms of the angle difference between each FA normal and the direction θ_model_, from cells without treatment (*A*), with calA (*B*), and with blebbistatin (*C*). (*D*) Average FA densities among the cells under six different cases (the error bars represent the standard deviation). Statistical differences were found between strain and control experiments for all 3 treatments, as well as between treatments (unpaired t-tests, p < 0.05), except between calA and DMSO (unpaired t-tests, p > 0.05). The number of FAs analysed in each histogram is n_strain_=7568 (61 cells) and n_control_=2607 (35 cells) for the drug-free condition (A, D), n_strain_=5164 (71 cells) and n_control_=4119 (44 cells) for the calA condition (B, D), and finally n_strain_=1904 (62 cells) and n_control_=889 (29 cells) for the blebbistatin condition (C, D). Table S2 of the ESI† specifies the number of FAs analysed in each case for both the control and stretched experiments. (*E*-*F*) Immunofluorescence images of a HFF cell following 11 hours of cyclic stretching along the x-axis (1 Hz) on the fibronectin-coated PDMS membrane embedded in the microdevice. Immediately after the stretching process, the cells were fixed and stained for actin filaments (red) (*E*, *G*), vinculin (green) (*F*, *G*), and DNA (blue) (*G*). The images from the laser scanning confocal microscope were post-processed with ImageJ to perform the z-projection, background removal, and contrast enhancement. The scale bar is 25 μm.

## Discussion

### Cells avoid large positive axial strains

Our results show that in the region of constant strain amplitude and direction (central region, see ESI† Fig. S3), the cells reoriented approximately perpendicularly to the stretching direction, which is in general agreement with a large body of work performed mainly with uniform strain amplitude devices^10–18^. Theoretical models have been proposed to describe the reorientation response of cells to cyclic stretching. For high frequency stretching (∼1 Hz) and moderate strain amplitudes (∼5-15 %), different experiments and models suggest several possible preferential orientations. Various studies corroborate models in which the preferential cell orientation is the minimum absolute strain direction, i.e. the direction minimizing length changes along the major cell body axis^10,17,19^. Building upon previous work^18^, Kaunas *et al.* proposed a model with this predicted preferential orientation, based on the premise that the disassembly rate of the stress fibers should be the smallest along this direction^47^. Considering a simplified model where FAs are treated as catch bonds destabilized by cyclic stretching, Chen *et al*. arrived at similar conclusions with respect to the preferential orientation^24^. On the other hand, Safran and De developed a comprehensive model based on effective free energy considerations and in which cells are treated as force dipoles^22^. Their model is adaptable to allow for two possible preferential orientations: minimum absolute strain direction or minimum stress direction. In the latter case, the corresponding preferential orientation is perpendicular to the maximum principal strain direction, rather than along the minimal absolute strain direction. Based on empirical observations, Neidlinger-Wilke et al. had also proposed previously that cells should preferentially reorient along this direction^12^ (even if this requires undergoing axial compression), and several experimental studies corroborate such alternative predictions^14–17^. Finally, Livne et al. recently proposed a model based on dissipative relaxation of stored elastic energy which was highly successful in explaining their experimental results^20^. Importantly, since uniaxial stretching of a deformable membrane generally results in orthogonal compression, the non-zero Poisson ratio greatly impacts the predicted preferential orientation in many of the models noted above. In our case, the membrane’s Poisson ratio was approximately 0.5 and so was the actual ratio of X-compression over Y-stretching in the central portion of the membrane. The resulting minimum absolute strain direction (zero-strain direction) is 35° with respect to the x-axis, which is not the preferential orientation we observed (see Fig. S3 of the ESI†). The preferential orientation predicted by the model of Livne et al. is ∼30°, which is also markedly different from what we observed. We rather observed that the fraction of cells subjected to an approximately uniform strain field in our device preferentially oriented themselves perpendicularly (0°) to the maximum principal strain direction. Interpreted from the point of view of the model by Safran and De^22^, our results thus suggest that HFF cells stretched in our device preferentially align along the minimal stress direction rather than that of the zero strain. The lack of consensus concerning the exact preferential orientation under uniform strain field suggests a dependence on the cell type and experimental conditions. Finally, we note that in general, the broadness of the observed orientation distributions in cyclic stretching experiments can be attributed to stochastic contributions associated with the statistical nature of the chemical reactions involved in the mechanosensing process and variations in the cell population^22,48^.

The extent of cell alignment in our experiments was found to increase with the amplitude of the strain (Fig. 3A), as reported previously for human fibroblasts^12,48^ and for other cell types^9,11,49^. A feature of this relationship identified in previous studies is the presence of a strain amplitude threshold, under which no significant alignment is observed. It was reported to be ∼4 % for human fibroblasts^12^. This behaviour is in qualitative agreement with our data, although Fig. 3A suggests a slightly lower threshold. A more precise threshold analysis would not be relevant here because of the strain gradient interplay.

### Cells seek to avoid strain gradient

The cell reorientation behaviour in a non-uniform strain field has been investigated previously, but other studies failed to show a dependence upon the strain gradient. This can be explained by the low gradient amplitudes applied, the gradient direction being systematically along that of the strain, or the absence of comprehensive analyses of cellular reorientation with respect to the strain gradient field. In our case, the successful observation of this effect was made possible by the combination of biologically relevant strain gradient amplitudes (0 to 14% mm^-1^) and the complex strain field geometry at a scale similar to cell sizes in which the strain and the gradient directions are decoupled. Importantly, as shown above (Fig. 1A, 2D, and 3B), in regions of large gradient amplitudes, cells tended to align perpendicularly to the gradient direction. This observation suggests that cells respond to the gradient to avoid being subjected to different strain amplitudes throughout its different adhesion sites. This apparent avoidance of the strain gradient is analogous to the previously established avoidance of length increases (or changes) during cyclic stretching. Interestingly, cell reorientation is dependent on the strain gradient amplitude and the strain amplitude, respectively shown in Fig. 3A and 3B. These relationships are reflected in the choice of the simple empirical model.

It is worth mentioning previously reported observations enabled by different devices producing non-uniform strain fields. In particular, the use of uniaxial devices allowed the generation of membrane strain gradients, which were used to demonstrate the influence of the strain amplitude under static stretching^34,38^. However, in these cases, the strain gradient was oriented along the stretching direction, preventing investigation of its influence on cell orientation. In other instances, radially deformable membranes producing radial strain gradients showed that under cyclic stretching, cells orient themselves perpendicularly to the principal strain direction^33^ and the alteration of mRNA expression is strain amplitude dependent^36,37^. Interestingly, while no gradient-dependent reorientation was observed, likely due to the device’s radial symmetry, gradient-dependent mRNA expression was observed (which is compatible with our results). Also relevant to the present study, Morita *et al.* studied human bone marrow mesenchymal stem cell reorientation in a non-uniform cyclic strain field^9^. The strain gradient was on the order of 0.5 %mm^-1^, and it was mainly in the stretching direction. They methodically analysed the potential dependence of the cell reorientation on the strain gradient, but found none under their conditions. Finally, Ohashi *et al*. employed a cyclically stretched membrane exhibiting a very large gradient (200 %mm^-1^) in the stretching direction to study stress fiber formation^35^. Interestingly, it was found that variations in the local development of intracellular stress fibers are associated with variations in applied strain amplitude underneath this cell (local response).

It must be noted that the cell alignment response to strain gradient found in our study is distinct by its non-local manifestation. In general, the capacity for a body to sense a strain gradient requires it to spatially locate and compare different strain amplitudes. Although speculative, we hypothesize that the global cellular response to strain gradient implies that cells are be able to pinpoint this information at each adhesion site, and coordinate these signals (strain amplitude, strain direction, and location under the cell) to determine optimal alignment. In this context, the cell exhibits a coordinated and integrated global response that is partially based on the spatial variation in strain amplitudes sensed locally in different cell areas. This is particularly easy to observe in our device because at the scale of a single cell, both the gradient and strain directions are relatively constant, only the strain amplitude varies significantly.

### Contractility is central in the cyclic stretched-induced cell alignment

FA analysis and drug treatment experiments were also performed in this study. The prestress condition and actomyosin contractility are directly involved in cell reorientation^21,27,50^. Therefore, reagents affecting the actomyosin cytoskeleton are most likely to strongly impact stretch-induced cellular reorientation^26^. Two such reagents are calA and blebbistatin, which promote^45^ and inhibit^27,46^ contractility through the modulation of myosin-II activity, respectively. Our observation is similar to that of Zhao *et al.*^27^ who reported a high degree of perpendicular reorientation among the calA-treated fibroblast cell population. In their case, this response was faster than without the pharmacological treatment, but their trends converged after several hours. In our case, after 11 hours of stretching, the reorientation responses of the calA-treated cells and the untreated cells were not significantly different. It was also previously reported that a sufficient concentration of blebbistatin largely supresses the reorientation of 3T3 fibroblasts cells and stress fibers under cyclic stretching^26,27^. This behaviour, involving stress fiber tension reduction, is in qualitative agreement with our results, which supports the notion that contractility is central to cellular reorientation.

### The reorientation of FAs is primarily driven by internal forces under our conditions

The cell machinery enabling the sensing and transduction of extracellular mechanical cues comprises not only the cytoskeletal network but also the FAs; the two are intrinsically connected^29,31^. The investigation of FA density and orientation under cyclic stretching, following a change in myosin-II activity, can provide insight on their roles in cellular reorientation. Our results show that the FAs mostly reorient perpendicularly to the strain direction upon cyclic stretching, in agreement with previously reported studies^23,30,51^. As well, we quantitatively show that FAs align parallel to the actin stress fibers, since the later mostly follow cell orientation, as expected^20,52^. On a broader level, it was proposed that the FAs elongate by growing in the direction of a pulling force^53^. This suggests that in our experiment, the primary forces driving FA reorganization are internal (rather than substrate stretch); i.e. they are mainly associated with actin assembly and actomyosin contraction.

### Contractility affects FA density and reorientation behaviour

We found that under cyclic stretching, the inhibition of cell contractility via blebbistatin treatment resulted in the inhibition of FA alignment and a significant reduction in FA density. The blebbistatin-induced loss of FA density is in agreement with a previously reported study^29^. Our results further suggest the strong dependence of the FA fate on the actin-induced forces. Moreover, we observe a small – but statistically significant – effect of the substrate strain itself on the FA density following 11 hours of cyclic stretching (for all three contractility conditions). In contrast, a larger increase in FA density, induced by stretching, has been reported for other cell types when studied on different time scales^54–56^. Furthermore, calA (without stretching) was previously shown to increase FA density but only on a short time scale^57^. This is in contrast with the absence of significant difference in FA density observed here between the calA-treated and non-treated cells, when studied over our much longer time scale (11 hours). Overall, our results highlight the fact that FAs are not only implicated in the mechanosensing process, but their behaviour is also an integral part of the mechanoresponse.

### Potential mechanisms involved in strain gradient sensing

Forces acting on the ECM-integrin-cytoskeleton complex are believed to be sensed via numerous mechanisms^58^. In the context of uniform strains, it has been shown that forces exerted by the ECM on the cell can promote cytoskeleton and focal adhesion assembly and reorganization. The exact mechanisms for tension-mediated FA maturation upon mechanical stimulation are not fully understood, but they are known to involve diverse protein recruitment and integrin clustering^59^. A specific example may include the reinforcement of the integrin-actin linkage by unfolding specific talin domains via stretching, thus allowing vinculin binding^60^. This mechanism was recently revealed to be involved in force transmission and transduction in the context of substrate stiffness sensing through the actin-talin-integrin-fibronectin clutch^61,62^. Stabilization of integrin adhesions through the vinculin recruitment induced by myosin-dependent tension has also been shown via FAK and SRC-mediated phosphorylation of paxillin^59^. More recently, in fibronectin-based microenvironments, mechanosensing is believed to involve α_V_-class integrins which were shown to play a role in the structural cell adaptation to forces^63^. Interestingly, the expression of α_V_β_3_ was specifically shown to be upregulated in cyclically stretched in endothelial cells^56^. Moreover, α_5_β_1_ was demonstrated to enable force generation and the cooperation of the latter with α_V_-class integrins was shown to enable rigidity sensing^63^. We note that the mechanosensing machinery enabling strain sensing bears similarities with that associated with stiffness sensing. Importantly, in both cases, stiffness- or stretching-induced signal transduction cascades activate the RHO-ROCK pathways which in turns regulate the myosin light chain phosphorylation and thus enable tension and traction force generation via the actomyosin cytoskeleton^58^.

Through the above molecular mechanisms, cells are able to probe and respond to the spatial profile of the ECM stiffness, possibly at the scale of the individual FAs^64^ and thus with micron precision. This enables cells to sense stiffness gradient and undergo gradient-guided migration (durotaxis)^44,65^. In a similar way, it was previously shown that cells are sensitive to the underlying strain profile at the sub-cellular scale^35^, and thus again possibly at the scale of individual FAs. Although our data suggest that the preferential alignment of the FAs is affected by both the strain and the strain gradient properties of the ECM, it is difficult to know if the latter behaviour results from the direct guidance of the ECM external force onto the FAs individually, or if it rather results from the action of the internal cytoskeleton tension adapting to the external strain gradient.

We speculate that the exact underlying mechanisms that ultimately govern strain-gradient sensing and avoidance are likely highly similar to the pathways identified above. However, we note that these pathways are highly dependent on the cell type, ECM and stretching conditions. Therefore, it is currently beyond the scope of this study to attempt to generalize a specific mechanism. Following the logic of Tse and Engler^65^ in the context of stiffness gradient sensing, we suggest that the strain gradient sensing process involves the creation intracellular signalling gradients such as gradients of RHO-ROCK activity. Another possibility would be a gradient of talin-mediated force transmission resulting from spatial variations in the duty cycles of vinculin-binding site exposition across the cell, directly correlating with different stretching amplitudes. We hypothesize that such signalling gradients, together with direct variation of force imposed across the cell, result in increased cytoskeleton tension gradients within the cell. This will be explored in future work utilizing the framework we have now presented and established in this study. It is well established that cells seek to maintain mechanical homeostasis under changing ECM properties^58^. Therefore, gradient avoidance suggests that cell homeostasis also includes a certain degree of spatial tension uniformity across the cytoskeleton, possibly at the expense of deviating from the preferential orientation determined by the average strain direction.

## Materials and methods

### Microdevice geometry and fabrication

The cells were immobilized on a thin suspended PDMS (Sylgard184, Ellsworth Adhesives Canada Corporation, Stoney Creek ON, Canada) membrane and cyclically stretched within microdevices which have a modified design similar to the one we previously reported^13^. The chip size and geometry allow for high strain gradient amplitudes (from 0 to 14%mm^-1^) to be generated on the membrane, while keeping the strain amplitude relatively low (from 2 to 10%, see Fig. 1A, B). The unique strain field pattern was achieved by stretching a 1.6 mm x 1.6 mm x 10 μm membrane with four fixed boundaries, using 4 vacuum chambers, all embedded in the PDMS chip. The action of the vacuum chambers deforms the cell chamber side walls (120 μm thick) in which the membrane is anchored, allowing the stretching motions. The device comprises three layers consisting in a suspended membrane inserted between the micro-patterned top and bottom parts. The membrane’s stiffness is estimated to 0.5 MPa based on previously characterized PDMS substrates prepared under similar conditions to ours^66,67^. Additional fabrication details are provided in the ESI† (see Fig. S1).

### Membrane strain field calculation

The non-uniform nature of the strain field requires a detailed point-by-point characterization to accurately analyse the cell reorientation based on the local field applied to each cell. Fluorescent beads (FluoSpheres, 200 nm, Invitrogen, CA, USA) were incorporated in the floating 10 μm thick membrane for precise strain field characterization. The full stretching motion of the substrate was divided in 10 frames (by applying different pressure in the vacuum chambers) and imaged with an EPI fluorescence microscope (Nikon, <u>Tokyo, Japan</u>). The beads displacements were tracked with a homemade Matlab program in which the Green strain matrix elements were subsequently calculated for each initial bead position. From these, two principal strain directions and amplitudes can be extracted. In such coordinate systems, the shear strain is zero. First, the maximum principal strain **ε_1_**(x,y) is associated with the direction of maximum stretch at a given membrane position. Second, the minimum principal strain **ε_2_**(x,y) (with direction that is orthogonal to that of **ε_1_**(x,y)), is associated with the direction of minimum stretch (or maximal compression in the case of negative **ε_2_**). Note that **ε_1_**(x,y) is of primary interest here since it largely dominates **ε_2_**(x,y) across the membrane. The gradient of ε_1_(x, y) was then calculated to produce the gradient amplitude |∇ε_1_(x, y)| and gradient angle θ_gradient_(x, y) maps (see Fig. 1 b). Additional details on the strain calculations are provided in the ESI†.

### Experimental procedures

The assembled microdevices were air-plasma treated (Glow Research, Tempe, AZ, USA) for 5 minutes at 70 W prior to sterilization with 70% ethanol for 5 minutes. After flushing the ethanol with autoclaved phosphate buffered saline (PBS) solution, the membrane was functionalized for 4 hours at 37°C with a solution of fibronectin 10 μg/ml of HEPES-buffered salt solution (HBSS; 20 mM HEPES at pH 7.4, 120 mM NaCl, 5.3 mM KCl, 0.8 mM MgSO_4_, 1.8 mM CaCl_2_ and 11.1 mM glucose). A solution of suspended cells in the culture medium with a density of 5x10^4^ cells/ml was injected into the 5.2 μl device’s cell chamber. The devices containing the cells were left in a standard incubator for 15 hours to let the cells attach and spread on the fibronectin coated membrane. The well adhered cells were then cyclically stretched for 11 hours at 1 Hz by activating the vacuum pumps while keeping the microdevice in the incubator during the whole experiment. The stretching time of 11 hours was chosen to be well above the characteristic time for the strain-induced reorientation of fibroblasts under our conditions^10^ and the stretching frequency was chosen for its physiological relevance^68–70,8^. The maximum strain amplitude field varies from 2% to 10% across the membrane, as depicted in Fig. 1A. A homemade Labview program controlled the maximum amplitude and frequency of the sinusoidal stretching wave form.

### Cell preparation and drug treatment

HFF cells (American Type Culture Collection (ATCC), Manassas, VA, USA) were cultured in Dulbecco’s Modified Eagle’s Medium (DMEM) supplemented with 10% FBS and 1% streptomycin/penicillin (Hyclone Laboratories, Logan, UT, USA). All cells were maintained at 37°C and 5% CO_2_ in a standard incubator. The drugs (Sigma Aldrich, St. Louis, MO, USA) used were stored in dimethyl sulfoxide (DMSO) stock solutions and were added to the cells immediately before starting the stretching process. DMSO with the same final concentration (0.1%) was also added to the control experiments that were ran without pharmacological treatment. The drug treatments were applied to cells for the entire time of the experiments (11 hours). The final concentration of calA (2 nM) was based on previous studies^57,71,72^ and by considering the long duration of the treatment (11 hours). More specifically, it was selected to avoid cell detachment from the substrate and rapid morphology changes (rounding) caused by the increased myosin-II contraction^73,74^. The final blebbistatin concentration (10μM) was chosen to enable relevant comparisons with related studies ^30,72,75^.

### Cell fixing, immunofluorescence straining, and imaging

Immediately after the 11 hour stretching process, the cells were fixed with 3.5% paraformal-dehydeandpermeabilized and permeabilized with 0.5% TritonX-100. The cells enclosed in the microdevice were then kept in a cold PBS solution and imaged with a Nikon TiE inverted phase contrast EPI microscope with a long working distance 10x objective in order to acquire the cell orientation data. Actin filaments were then stained with Phalloidin conjugated to Alexa Fluor 546 (Invitrogen) and the DNA with DAPI (Invitrogen). A monoclonal mouse anti-vinculin antibody and a rabbit anti-mouse IgG secondary antibody conjugated to Alexa Fluor 488 (Invitrogen) were used to stain vinculin. The details of the immunofluorescence staining protocols have been published previously^76,77^. These protocols have been adapted to suit the microdevice environment (see ESI† for details). An upright laser scanning multiphoton confocal microscope (Nikon **A1RsiMP**) with a long working distance 25x objective (NA=1.1) was employed to image the actin fibers, the nuclei, and the vinculin proteins.

### Cell orientation and FA analysis

The phase contrast images of the cells and the fluorescent images of the FAs were post-processed in ImageJ. The fluorescence images were first z-projected. To correct for background noise, a Fast Fourier Transform bandpass filter was applied to the phase contrast images, while the subtract background tool was employed for the fluorescent images. In both cases, binary images were then obtained from the adaptive threshold plugin which optimally captures the cell and FA morphologies despite the non-homogenous images intensities. Finally, the analyze particles tool was used for the extraction of the cell and FA positions, sizes, and orientations.

A homemade Matlab program was developed to carry out the analysis. The cells and FAs located within 8% of the edges of the devices were excluded to discard edge effects. The orientation of each cell (or FA) was compared to the local strain parameters. More precisely, we computed the angle difference between the cell (or FA) normal and either the maximum principal strain angle or the model angle. The equation of the latter, which is presented in the Results section, includes the weighting parameter β. All the analysis was performed with the same β value after its initial optimization with the data set of Fig. 2D. To construct the histograms, the angles were converted to a 0 to 90° range.

### Conclusion

In summary, cyclic stretching experiments under diverse conditions were performed in micro-fabricated devices exhibiting a unique highly non-uniform strain fields. These experiments revealed that fibroblast cells seek to avoid strain gradients similarly to the way they avoid stretching. The degrees of these avoidances are shown to depend on the strain gradient amplitude and the strain amplitude respectively. Importantly, by identifying the strain gradient as a subtle mechanical cue, our results demonstrate that cells can integrate diverse physical signals simultaneously and exhibit a coordinated response. Furthermore, we demonstrated that under non-uniform strain field, human fibroblast FAs also reorient in a similar fashion to the cells, and that their density is affected by stretching forces. Finally, myosin-II contractility was shown to impact FA and cell reorientations, since its inhibition prevented the alignments of both and yielded a reduced FA density.

Our work reveals that strain gradients are mechanical cues capable of influencing and modulating global cellular organization and orientation. This opens a new line of investigation regarding the role of the strain gradient in regulating certain cell functions and how this regulation can be compromised by cell or cell matrix defects. This study paves the way to a better understanding of certain in vivo pathologies which may lead to new pharmaceutical approaches, in addition to providing new insights on the fundamental interactions between the cells and their microenvironment.

## Acknowledgements

This work was supported by individual Natural Sciences and Engineering Research Council (NSERC) Discovery Grants to M.G. and A.E.P. S.C-L. was supported by NSERC Postgraduate Scholarships-Doctoral (PGS D). M.G. acknowledges support from the Ontario Ministry of Research and Innovation, and the Canada Foundation for Innovation (CFI). A.E.P. gratefully acknowledges generous support from the Canada Research Chairs (CRC) program.

